# Non-Invasive classification of macrophage polarisation by 2P-FLIM and machine learning

**DOI:** 10.1101/2022.01.22.477332

**Authors:** Nuno G.B. Neto, Sinead A. O’Rourke, Mimi Zhang, Hannah K. Fitzgerald, Aisling Dunne, Michael G. Monaghan

## Abstract

In this study, fluorescence lifetime imaging of NAD(P)H-based cellular autofluorescence is applied as a non-invasive modality to classify two contrasting states of human macrophages by proxy of their governing metabolic state. Macrophages were obtained from human blood-circulating monocytes, polarised using established treatments, and metabolically challenged using small molecules to validate their responding metabolic actions in extracellular acidification and oxygen consumption. Fluorescence lifetime imaging microscopy (FLIM) quantified variations in NAD(P)H-derived fluorescent lifetimes in large field-of-view images of individual polarised macrophages also challenged, in real-time with small molecule perturbations of metabolism during imaging. We uncover FLIM parameters that are pronounced under the action of carbonyl cyanide-p-trifluoromethoxyphenylhydrazone (FCCP) which strongly stratifies the phenotype of polarised human macrophages. This stratification and parameters emanating from a FLIM approach, served as the basis for machine learning models. Applying a random forest model, identified three strongly governing FLIM parameters, achieving a ROC AUC value of 0.944 when classifying human macrophages.

## 2. Introduction

Two-photon fluorescence lifetime imaging microscopy (2P-FLIM) is a non-destructive modality that can interrogate exogenous and endogenous fluorophores, while providing high spatial and temporal resolution to image 2D and 3D cell cultures and biopsies *in vitro* and *in vivo* (Neto et al. 2020, Okkelman et al. 2019, Skala et al. 2007). Reduced nicotinamide adenine dinucleotide (NAD(P)H) and flavin adenine dinucleotide (FAD^+^) can be studied and quantified using 2P-FLIM, providing insight into cellular metabolism (Lakowicz et al. 1992, Neto et al. 2020, Skala et al. 2007). The average time for a fluorophore to return to the ground state from the excited state while emitting fluorescence is known as the fluorescence lifetime and offers extended insight on NAD(P)H protein/enzyme binding (Lakowicz et al. 1992). NAD(P)H is noted as having a long fluorescent lifetime when enzyme-bound and a short fluorescence lifetime when free in the cytoplasm with 2P-FLIM facilitating quantification of these fluorescence lifetimes and their respective proportions. This information has been harnessed to spatially assess the metabolism of murine tumour tissues, murine-derived organoid cultures, human T-cells, human B-cells, murine macrophages and cultures subjected to bioreactor engineered nutrient gradients in which fluorescence intensity and lifetime of NAD(P)H and FAD^+^ were influenced by microenvironmental metabolites (Alfonso-Garcia et al. 2016, Floudas et al. 2020, Okkelman et al. 2019, Perottoni et al. 2021, Skala et al. 2007, Szulczewski et al. 2016, Walsh et al. 2020). NADH and NADPH fluorescence properties are identical, hence NAD(P)H refers to these intracellular pools combined (Huang et al. 2002). Due to its microscopy-based spatial approach, 2P-FLIM can be applied to limited cell numbers which is beneficial in assessing limited sample numbers or validating metabolism-based therapies (Peterson et al. 2018, Shields et al. 2020).

Macrophages are essential components of the host immune response and key regulators of homeostatic function. In addition to host defence, they are intimately involved in tissue homeostasis and play a key role in pathologies including heart failure, diabetes and cancer (Mosser et al. 2008). Macrophages adopt specific polarisation states for different functions and mechanisms of action which exist in a spectrum, on which classically activated (M1) and alternatively activated (M2) macrophages occupy opposite ends (Gordon 2003, Mosser et al. 2008). In vitro, M1-macrophage behaviour can be evoked using IFNγ and/or microbial products such as LPS with their phenotype described according to their secretion of significant amounts of proinflammatory cytokines such as TNFα and IL-β, enhanced endocytic and ability to kill intracellular pathogens (Adams 1989, Martinez et al. 2008). M2-macrophage behaviour exists more heterogeneously and can be triggered by IL-4 or IL-13, IL1-β, TGF-β, IL-6, phagocytosis of apoptotic cells or association with a tumour microenvironment, generating M2a, M2b, M2c, M2f and tumour-associated (TAM) macrophages, respectively(Graney et al. 2020, Mantovani et al. 2017, Mantovani et al. 2002). M2 macrophages exhibit decreased expression of protein membrane markers such as CD14 and CCR5, and increased fibronectin-1 production, and attribute to the downregulation of pro-inflammatory cytokines (Gratchev et al. 2001, Martinez et al. 2008, Wang et al. 1998). Most often macrophage behaviour is assessed using end-point assays using cytokine measurements, gene analysis and staining of surface markers but there is a gathering shift towards non-invasive modalities to speed up this process and obtain spatio-temporal analysis. Two recent examples of this include the use of raman microscopy to map the lipidomic spatial signature of polarised macrophages (Feuerer et al. 2021), and the use of average fluorescent lifetime parameters to discern phenotype (Kröger et al. 2021).

Here we use an imaging-based approach primarily focusing on metabolic machinery characteristically employed by polarised human macrophages, as they pose huge potential in the next-generation of therapeutics for inflammatory disease. Human macrophage function and their metabolism are inextricably linked (Van den Bossche et al. 2017). IFNγ activated human macrophages (IFNγ-M1) are primed for enhanced inflammatory responses by stabilising HIF1α levels, activation of the JAK-STAT pathway, and increased production of IL-1β, all of which are dependent on enhanced levels of glycolysis (Wang et al. 2018). Alternatively activated human anti-inflammatory macrophages, most often observed *in vitro* through stimulation with IL-4, (IL-4-M2) are defined by an intact tricarboxylic acid cycle (TCA), enhanced OxPhos, increased fatty acid synthesis (FAS) and fatty acid oxidation (FAO) (O’Neill et al. 2016, Van den Bossche et al. 2017). In a nutshell, IFNγ-M1 macrophages are more active in glycolysis, while IL-4-M2 macrophages are more dependent on oxidative phosphorylation for their energy production. When assessing metabolism and bioenergetics, most methods are based on extracellular flux assays, measurement of cellular oxygen consumption, exogenous staining, radio-labelling nutrients and gas chromatography – mass spectrometry (GC-MS), requiring a substantial amount of sample processing yet still poses limitations due to short lived oxidative metabolites (Fall et al. 2020, Koo et al. 2016, Ma et al. 2020, Vivekanandan-Giri et al. 2011).

2P-FLIM imaging of NAD(P)H provides five fluorescence lifetime variables (τ_1_, τ_2_, α_1_, α_2_, τ_avg_) and one fluorescence intensity-based variable (ORR) which can be applied to understand the polarisation linked-metabolic state of IFNγ-M1 and IL-4-M2 macrophage phenotypes. Furthermore, additional information can be obtained from these measurements by real-time perturbation of basal metabolism and metabolic capacity through sequential presentation of excess nutrients (glucose), knockdown of intercellular pathways via metabolic inhibitors and uncomplexing of mitochondrial coupling which yields an extended array of information. Considering such a vast array of information, we utilised Uniform Manifold Approximate and Projection (UMAP), instead of principle component analysis (PCA) or t-distributed Stochastic Neighbor Embedding (t-SNE) due to its processing speed, capability to preserve the global and local structure of the data and ability to use non-metric distance functions(McInnes et al. 2018). To aid this knowledge extraction, machine learning algorithms are employed for supervised pattern recognition in large data-sets. Machine learning can be defined as a computational and data science methodology which uses human-labelled experimental information to make accurate predictions or improve performance (Mohri 2012). Machine learning algorithms have been used to characterise DNA- and RNA-binding proteins, determine genetic and epigenetic contributions of antibody repertoire diversity and to classify chronic periodontitis patients based on their immune cell response to ex-vivo stimulation with ligands(Alipanahi et al. 2015, Bolland et al. 2016, Culos et al. 2020). Machine learning algorithms are used to train classification models that allows the segregation of samples into different classes (for example in this study IFNy-M1 vs IL-4-M2, based on 2P-FLIM variables) whilst determining which variable is the most important for this task (Touw et al. 2013). For our work, we employed the Random Forest algorithm for pattern recognition and data segregation due to it high prediction accuracy and allowing to obtain information regarding variable importance for classification (Breiman 2001, Verikas et al. 2011). To successfully train the Random Forest classification algorithm, a k-fold cross-validation procedure was employed to reduced biases in the training data subset. Here, the raw data in randomly divided in to k subsets in which part of the data is used for training and the remaining for testing of the machine learning model, multiple times (Stone 1974). Afterwards, the Random Forest algorithm variables, such as the number of trees (*ntree*) and number of variables selected for the best split at each node (*mtry*) are optimised in order to reduce the out-of-bag (OOB) error, therefore improving the accuracy of the algorithm(Touw et al. 2013) (supplementary table 1). The optimised random forest machine learning model can be used to distinguish between the different populations of macrophages as well to measure which 2P-FLIM variables are the most important for this differentiation. To evaluate the predictive ability and efficiency of the data classification we calculated the area under the receiver operating characteristics curve (ROC-AUC), in which the specificity and sensitivity of predictive algorithm are plotted. This value summarises the predictive capabilities of the machine learning model from a value of 0 to 1, in which 1 is a perfectly accurate test. ROC-AUC values of 0.5 suggests no discrimination, 0.7 to 0.8 is considered acceptable, 0.8 to 0.9 is considered excellent and more than 0.9 is considered outstanding(Mandrekar 2010). Similar applications of machine learning have been explored by Walsh et al for classification of activated t-cells and Qian et al for quality control of cardiomyocyte differentiation (Qian et al. 2021, Walsh et al. 2020).

We hypothesise that 2P-FLIM of NAD(P)H and FAD^+^ provides quantitative information to evaluate and identify macrophage polarisation by proxy of their metabolism, with strong clinical potential as we use human cells from multiple donors in this study and the emergence of metabolic approaches to treat disease and inflammation. To test our hypothesis, we derived human macrophages from blood-circulating monocytes and polarised them into IFNγ-M1 or IL-4-M2 macrophages. We confirmed their cytokine and gene expression-related polarisation, while their metabolic behaviour was assessed via traditional extracellular flux assays. Finally 2P-FLIM was applied during which the real-time response of photonic variables to metabolism-challenging small molecules were measured. This study establishes 2P-FLIM as a mode to discriminate IFNγ-M1 and IL-4-M2 macrophages, which is most pronounced by their differential responses when treated with FCCP. This allows one to accurately classify macrophage polarisation by machine learning methods, e.g., the random forest model, and to evaluate single-cell heterogeneity. Our work reports, for the first time; a combination of dynamic real-time metabolic perturbations during FLIM analysis extensively studied using machine learning with macrophage polarisation as a reliable model system to demonstrate the impact and validity of this approach.

## 3. Materials and Methods

### Human blood monocyte-derived macrophage isolation

This study was approved by the research ethics committee of the School of Biochemistry and Immunology, Trinity College Dublin and was conducted in accordance with the Declaration of Helsinki. Leucocyte-enriched buffy coats from anonymous healthy donors were obtained with permission from the Irish Blood Transfusion Board (IBTS), St. James’s Hospital, Dublin. Donors provided informed written consent to the IBTS for their blood to be used for research purposes. PBMC were isolated and differentiated into macrophages as described previously (Mahon et al. 2020). The purity of CD14^+^CD11b^+^ macrophages was assessed by flow cytometry and was routinely >95%.

### Cytokine measurements

Macrophages (1 × 10^6^ cells/ml) were treated with IFNγ (20 ng/ml) or IL-4 (20 ng/ml) for 24 hours. Supernatants were harvested, and cytokine concentrations of TNFα and IL-10 were quantified by ELISA (eBioscience) according to the manufacturer’s protocol.

### Real-time PCR

Macrophages (1 × 10^6^ cells/ml) were treated with IFNγ (20 ng/ml) or IL-4 (20 ng/ml) for 24 hours. RNA was extracted using High-Pure RNA Isolation Kits (Roche) and assessed for concentration and purity using the NanoDrop 2000c – UV- Vis spectrophotometer. RNA was equalised and reverse transcribed using the Applied Biosystems High-Capacity cDNA reverse transcription kit. Real-Time PCR Detection System (Bio-Rad Laboratories, California) was used to detect mRNA expression of target genes. PCR reactions included iTaq Universal SYBR Green Supermix (Bio-rad Laboratories), cDNA Taqman fast universal PCR Master Mix and pre-designed TaqMan gene expression probes (Applied Biosystems) for CXCL9, CXCL10, MRC1, CCL13, and the housekeeping gene, 18S ribosomal RNA. The 2 ^− ΔΔCT^ method was used to analyse relative gene expression.

### Seahorse analyser

Macrophages were cultured at 1 × 10^6^ cells/ml for six days prior to re-seeding at 2 × 10^5^ cells/well in a Seahorse 96-well microplate and allowed to rest for five hours prior to stimulation with IFNγ (20 ng/ml) and IL-4 (20 ng/ml) for 24 hours. The Seahorse cartridge plate was hydrated with XF calibrant fluid and incubated in a non-CO_2_ incubator at 37°C for a minimum of eight hours prior to use. Thirty minutes prior to placement into the Seahorse XF/XFe analyser, cell culture medium was replaced with complete XF assay medium (Seahorse Biosciences, supplemented with 10 mM glucose, 1 mM sodium pyruvate, 2 mM L-glutamine, and pH adjusted to 7.4) and incubated in a non-CO_2_ incubator at 37°C. Blank wells (XF assay medium only) were prepared without cells for subtracting the background oxygen consumption rate (OCR) and extracellular acidification rate (ECAR) during analysis. Oligomycin (1 mM, Cayman Chemicals), carbonyl cyanide-p-trifluoromethoxyphenylhydrazone (FCCP) (1 mM, Santa Cruz biotechnology), rotenone (Rot) (500 nM), and antimycin A (AA) (500 nM) and 2-deoxy-D-glucose (2-DG) (25 mM, all Sigma-Aldrich) were prepared in XF assay medium and loaded into the appropriate injection ports on the cartridge plate and incubated for ten minutes in a non-CO2 incubator at 37°C. OCR and ECAR were measured over time with sequential injections of oligomycin, FCCP, rotenone and antimycin A and 2-DG. Analysis of results was performed using Wave software (Agilent Technologies). The rates of basal glycolysis, maximal glycolysis, basal respiration and maximal respiration were calculated as detailed in the manufacturer’s protocol and supplied in Supplementary Table 2.

### Two-photon fluorescence lifetime imaging microscopy (2P-FLIM)

2P-FLIM was performed on 24 hours-polarised macrophages seeded in *ibidi*® Luer μ-slides with a 0.8 mm channel height. 2P-FLIM was done using a custom upright (Olympus BX61WI) laser multiphoton microscopy system equipped with a pulsed (80 MHz) titanium:sapphire laser (Chameleon Ultra, Coherent®, USA), water-immersion 25× objective (Olympus, 1.05NA) and temperature-controlled stage at 37 °C. Two photon excitation of NAD(P)H and FAD^+^ fluorescence was performed at the excitation wavelength of 760 and 800 nm, respectively. Several studies have reported that two-photon excitation in the range of 720-760nm can be used to selectively excite NAD(P)H, while for FAD^+^ a excitation wavelength above 900nm is commonly used (Huang et al. 2002, Levitt et al. 2011).A 458/64 nm and 520/35 nm bandpass filter were used to isolate the NAD(P)H and FAD^+^ fluorescence emissions based on their emission spectra(Huang et al. 2002).

512 × 512 pixel images were acquired with a pixel dwell time of 3.81 μs and 30 s collection time. A PicoHarp 300 TCSPC system operating in the time-tagged mode coupled with a photomultiplier detector assembly (PMA) hybrid detector (PicoQuanT GmbH, Germany) was used for fluorescence decay measurements, yielding 256 time bins per pixel. TCSPC requires a defined “start”, provided by the electronics steering the laser pulse or a photodiode, and a defined “stop” signal, realized by detection with single-photon sensitive detectors. The measuring of this time delay is repeated many times to account for the statistical variance of the fluorophore’s emission. For more detailed information, the reader is referred elsewhere (Wahl et al. 2013). Fluorescence lifetime images with their associated decay curves for NAD(P)H were obtained, with a minimum of 1×10^6^ photons peak, and region of interest (ROI) analysis of the total cells present on the image was performed in order to remove any background artefact. The decay curved was generated and fitted with a double-exponential decay without including the instrument response function (IRF) (Equation (1)).

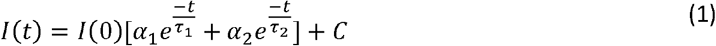

I(t) represents the fl uorescence intensity measured at time t after laser excitation; α _1_ and α_2_ represent the fraction of the overall signal proportion of a short and long component lifetime, respectively. τ_1_ and τ_2_ are the long and short lifetime components, respectively; C corresponds to background light. Chi-square statistical test was used to evaluate the goodness of multi-exponential fit to the raw fluorescence decay data. In this study, all of the fluorescence lifetime fitting values with χ^2^ < 1.3 were considered as ‘good’ fits. For NAD(P)H, the double exponential decay was used to differentiate between the protein-bound (τ_1_) and free (τ_2_) NAD(P)H. The average fluorescence lifetime was calculated using Equation (2).

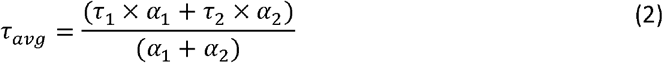

Intensity-based images of NAD(P)H and FAD^+^ were acquired, and their ratio was calculated using Equation (3) to obtain the optical redox ratio (ORR).

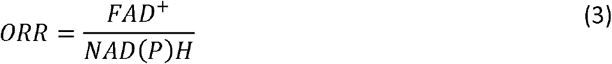

From the images acquired using 2P-FLIM, single-cell - analysis was performed using a custom-made script on Cell Profiler®(McQuin et al. 2018). The single-cell analysis was conducted in a similar way as the global 2P-FLIM analysis.

### Macrophage classification and machine learning

Uniform Manifold Approximate and Projection (UMAP) was used for dimension reduction, and z-score heatmaps were used to visualise the clustering of 2P-FLIM imaging datasets in global- and single-cell analysis(McInnes et al. 2018) UMAP was implemented in Python, and visualization in Graphpad). Random forest was applied to classify IFNγ-M1 and IL-4-M2 macrophages in both global- and single-cell approach. Random forest classification was implemented in R. Random forest hyperparameters include the number of decision trees in the forest, the number of features considered by each tree when splitting a node, and the maximal depth of each tree; the best hyperparameters were determined through grid search with the highest accuracy (supplementary Table 1). α_1_ variable was removed from the random forest model due to its deterministic relationship with α_2_ variable (supplementary Figure 1). We applied repeated 10-fold cross-validation to improve the estimated performance of the random forest model. Receiver-operating curves (ROC) were plotted, and areas under the curves (AUC) were calculated. In addition, random forest feature selection was employed to evaluate the weight of each 2P-FLIM variable to determine its relative importance in macrophage classification for both overall and single cell datasets. Support vector machine (SVM) and logistic regression models were compared with random forest model and yielded similar performance on the experimental data split to 80% training and 20% testing.

### Statistics

Each experiment was performed in at least four healthy donors (defined by N) with 3-4 technical replicates run for each experiment (defined by n), depending on the assay type. Normality tests were performed to determine the normal distribution of the data. For ELISA and PCR data, one-way ANOVA and Tukey’s test were used for comparing more than two groups. For Seahorse data, repeated-measures one way ANOVA was used to account for the variance in basal metabolism across donors. All statistical analysis was performed on GraphPad Prism™ 9.00 (GraphPad Software).

## 4. Results

### Macrophage polarisation with IFNγ and IL-4 induces metabolic reprograming

Human blood-derived macrophages were polarised by incubating in media containing IFNγ (M1) or IL-4 (M2) for 24 hours. Polarisation was confirmed using ELISA and RT-PCR. Cellular metabolic activity was analysed using a sequence of metabolic enzyme inhibitors and its effect measured by extracellular acidification ratio (ECAR), oxygen consumption ratio (OCR) and 2P-FLIM (Figure 1).

**Figure 1.**
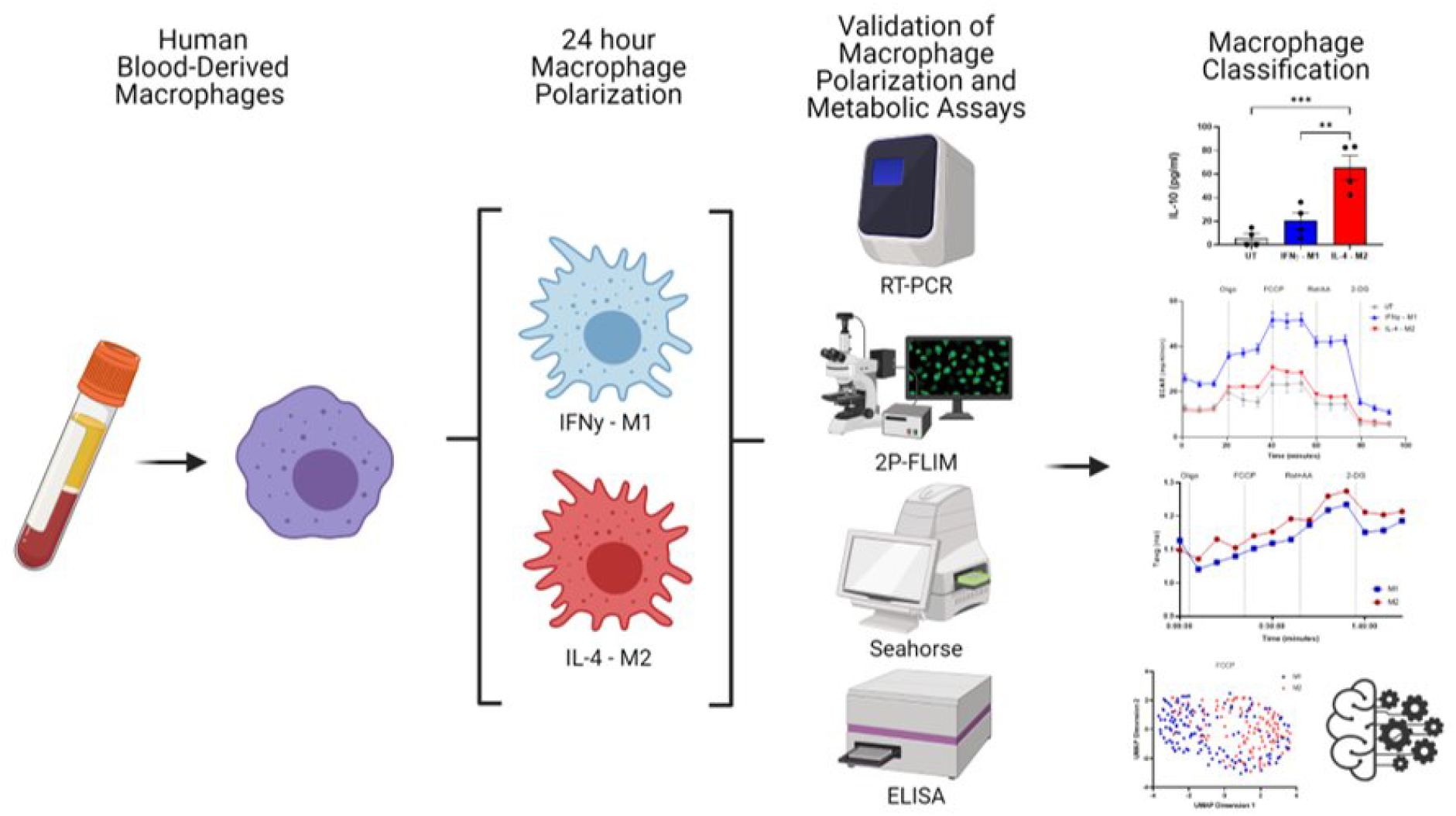
Overview of experimental work. Image created using biorender.com

A slightly higher amount of TNFα production was obtained for IFNγ-M1 when compared with IL-4-M2 macrophages. Regarding IL-10, a statistically higher production was measured in IL-4-M2 when compared with IFNγ-M1 and untreated macrophages (Figure 2A, D). For gene expression, a higher amount of CXCL9 and a statistically significant increase in CXCL10 in IFNγ-M1 macrophages were observed (Figure 2B, C). In addition, MRC1 and CCL13 were further expressed in IL-4-M2 macrophages when compared with untreated and IFNγ – M1 macrophages (Figure 2E, F).IFNγ-M1 macrophages have a higher dependence on aerobic glycolysis, whilst IL-4-M2 macrophages are more reliant on oxidative phosphorylation. We used extracellular acidification (ECAR) and oxygen consumption ratios (OCR) to certify this metabolic behaviour that is linked with macrophage polarisation. For ECAR and OCR, we used four different metabolic modulators in succession: oligomycin, FCCP, Rot/AA and 2-DG, to evaluate cellular metabolism (Figure 2).

**Figure 2.**
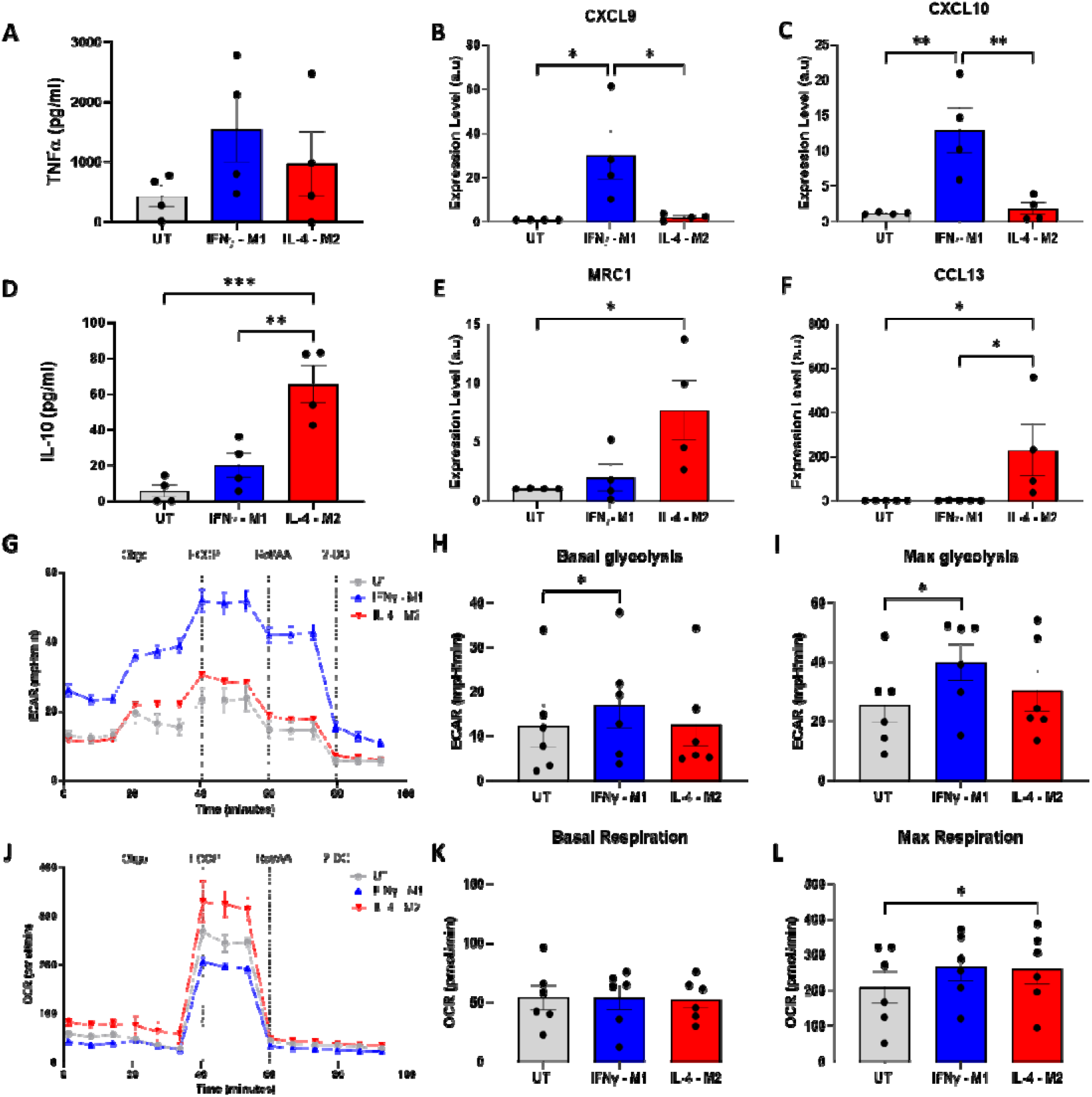
Validation of macrophage polarisation and metabolic profiling of IFNy-M1, IL-4-M2, and untreated (UT) macrophages. (A,B) ELISA of inflammatory cytokine TNFα and anti-inflammatory IL-10 in IFNy-M1, IL-4-M2, and untreated (UT) macrophages. (C,D,E,F) Evaluation of CXCL9, MRC1, CXCL10, and CCL13 gene expression in IFNy-M1, IL-4-M2, and untreated (UT) macrophages. (G,J) Extracellular acidification ratio (ECAR) and Oxygen consumption ratio (OCR) profile of IFNy-M1, IL-4-M2, and untreated (UT) macrophages when treated sequentially with oligomycin, FCCP, rotenone+antimycin A and 2-DG. (H,I,K,L) Area under the curve (AUC) values calculated from ECAR and OCR between each treatment. *p<0.05, **p<0.01, ***p<0.001.

IFNγ-M1 macrophages exhibited a higher ECAR and lower OCR in response to the treatments added, whilst IL-4-M2 and untreated macrophages had lower ECAR and higher OCR (Figure 2G, J). After obtaining the ECAR and OCR curves, the areas under the curves (AUC) were measured to reflect basal glycolysis, maximal glycolysis, basal respiration and maximal respiration.

IFNγ-M1 macrophages have a statistically significant increase of basal and max glycolysis when compared with untreated macrophages (Figure 2H, I). In addition, all macrophage types have similar basal respiration, whilst IL-4-M2 macrophages have a statistically significant increase in max respiration when compared with untreated macrophages (Figure 2K, L).

### 2P-FLIM captures metabolic shifts on IFNγ and IL-4 treated macrophages

2P-FLIM harvests NAD(P)H and FAD^+^ autofluorescence to infer cellular metabolism. NAD(P)H enzyme-bound state is characterised by a longer fluorescence lifetime whilst NAD(P)H free-state has a shorter fluorescence lifetime. NAD(P)H and FAD^+^ fluorescence intensity is measured in order to calculate the ORR (Equation 3). These fluorescence features enable the distinction between an OxPhos or glycolytic-dependent metabolism (Floudas et al. 2020, Neto et al. 2020, Okkelman et al. 2019, Perottoni et al. 2021, Schaefer et al. 2019, Skala et al. 1992, Walsh et al. 2020).

For this experiment, we seeded unpolarised macrophages in *ibidi*® Luer μ-slides in static conditions, and polarised the macrophages using IFNγ or IL-4 for 24 hours. These macrophages are derived from the same donors as per those presented in Figure 2. Afterwards, we followed the same subjection of metabolic enzymatic inhibitors applied in the ECAR/OCR experiment in which we treated the macrophages with oligomycin, FCCP, Rot/AA and 2-DG. During the time-course of the experiments the field of view was maintained so as to record single-cell metabolic variations (Figure 3A).

**Figure 3.**
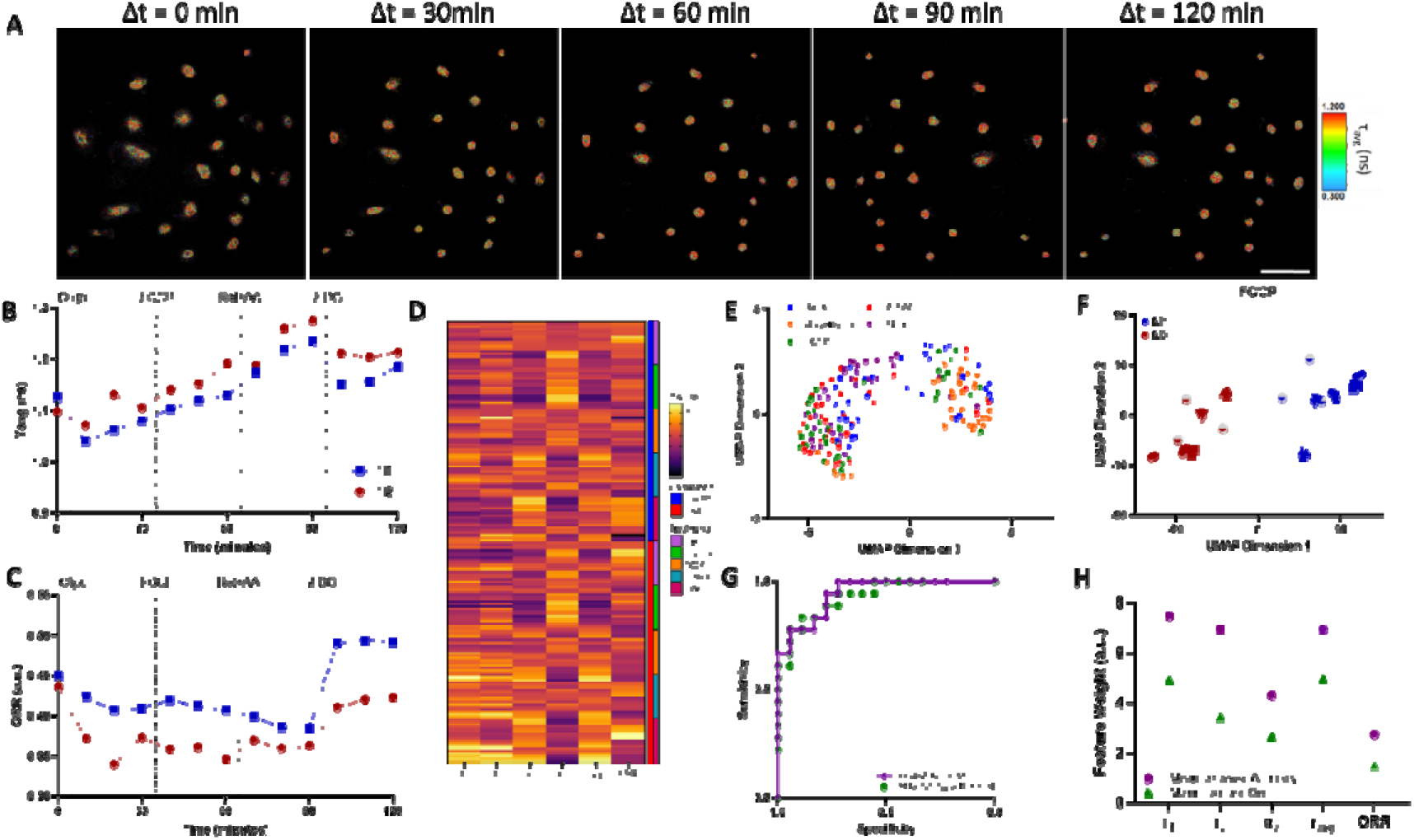
2P-FLIM Metabolimaging analysis. (A) Time-course imaging of representative (same field of view throughout) IFNy-M1 macrophages, scale bar:100μm (B,C) Average fluorescence lifetime (tavg) and optical redox ratio (ORR) values for IFNy-M1 and IL-4-M2 when treated sequentially with olygomycin, FCCP, rotenone+antimycin A and 2-DG of a representative donor. (D) z-score heatmap of 2P-FLIM acquired data for 6 donors separated by macrophage polarisation and metabolic inhibitor. (E) UMAP plot of 2P-FLIM variables after each treatment. (F) UMAP plot of 2P-FLIM variables after FCCP treatment. (G) Receiver-operator curve and area under curve values of random forest machine learning model applied to 2P-FLIM data after FCCP treatment. (H) 2P-FLIM weight features determined by mean decrease accuracy and mean decrease Gini of random forest model used to classify macrophages.

**Figure 4.**
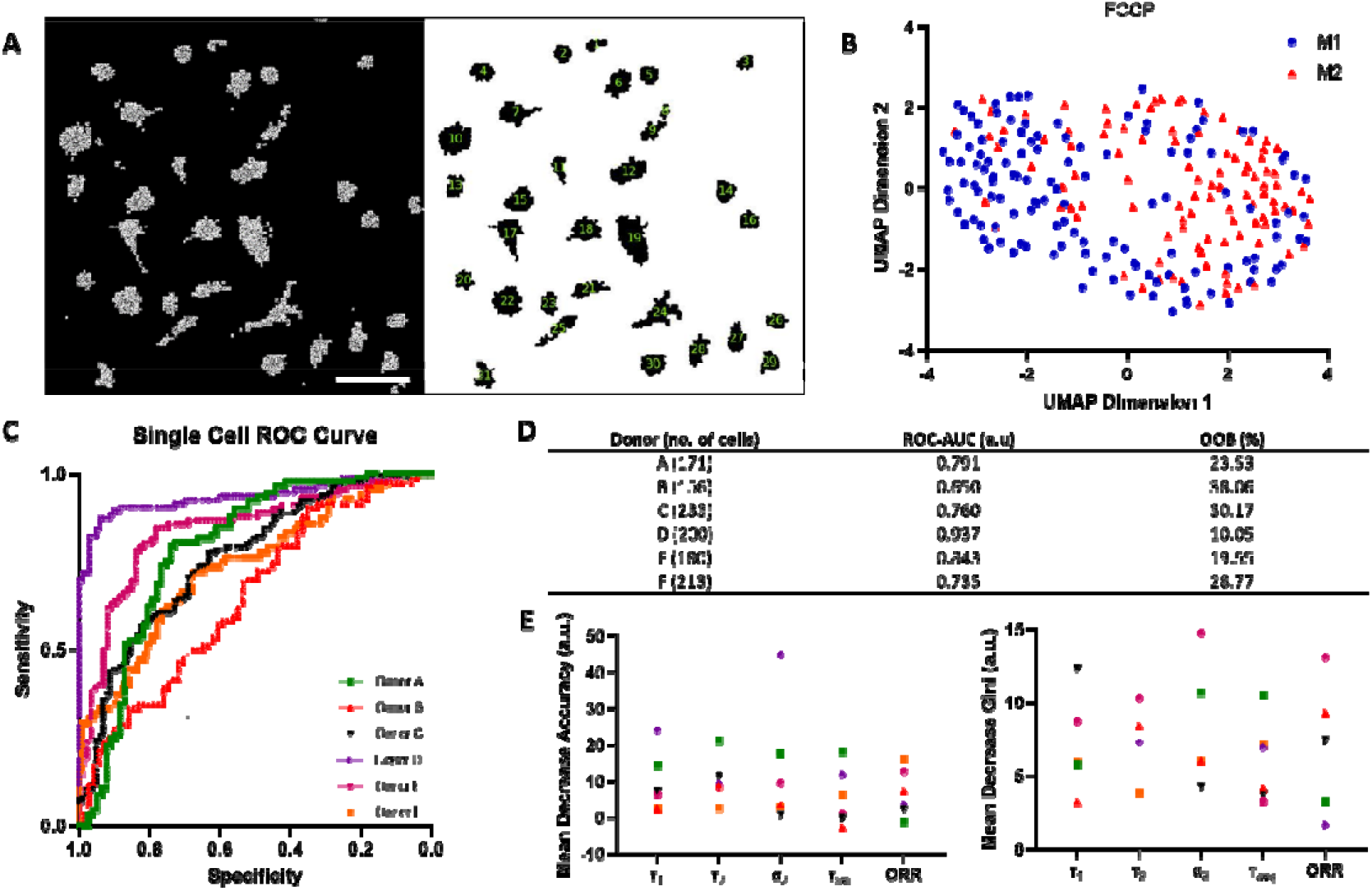
Single-cell 2P-FLIM imaging analysis. (A) Single-cell analysis using a custom-built Cell Profiler® script, scale bar=100μm. (B) Single-cell UMAP clustering of a representative donor after FCCP treatment using 2P-FLIM variables. (C) Receiver-operating curves (ROC) of random forest model for classification of macrophages of all human donors used in this study. (D) Area under the curve values for ROCs (ROC-AUC) and Out-of-bag error (OOB) calculated using random forest model for each individual donor. (E) Mean decrease in accuracy and mean decrease in Gini of each 2P-FLIM variable returned by the random forest model.

We derived the average fluorescence lifetime (τ_avg_) and optical redox ratio (ORR) of IFNγ-M1 and IL-4-M2 macrophages from the FLIM-data and observed an increasing trend of τ_avg_ in response to the application metabolic enzymatic inhibitors, with the exception of 2-DG in which a decrease of τ_avg_ is observed for both macrophage phenotypes (Figure 3B). With respect to ORR, there is a slight decreasing trend of ORR, followed by a raise in ORR with the 2-DG treatment for IFNγ-M1 macrophages. For IL-4-M2 macrophages, there is a decrease in ORR with the oligomycin treatment, followed by a consistent level, and finally an increase elicited by 2-DG (Figure 3C). We compiled all the 2P-FLIM variables: τ_1_, τ_2_, α_1_, α_2_, τ_avg_ and ORR into a representative z-score heatmap, stratified according to macrophage type and metabolic inhibitors across all donors. IFNγ-M1 macrophages have lower τ_1_, τ_2_, τ_avg_ and ORR values when compared with IL-4-M2 macrophages (Figure 3D).

### 2P-FLIM variables allow the classification of IFNγ-M1 and IL-4-M2 macrophages

Uniform Manifold Approximate and Projection (UMAP) was applied as a dimension reduction technique to FLIM variables associated with IFNγ-M1 and IL-4-M2 macrophages to visualise if this enables distinct clustering according to phenotype. The coordinates for each image were defined using a cosine distance function computed using the 2P-FLIM variables: τ_avg_, τ_1_, τ_2_, α_1_, α_2_ and ORR (Figure 3E). UMAP representation of the 2P-FLIM imaging data yielded two distinct clusters with the FCCP treatment providing a separation between IFNγ-M1 and IL-4-M2 macrophages (Figure 3E). If we consider the FCCP treatment alone, one can appreciate a clear segregation between IFNγ-M1 and IL-4-M2 macrophages across the six donors (Figure 3F). Based on this clear clustering observed with the FCCP treatment, classification models were established to classify macrophage polarisation from 2P-FLIM variables when treated with FCCP. Receiver operator characteristic (ROC) curves of test data reveal high accuracy for predicting macrophage polarisation in the full field of view using FCCP (AUC=0.944), when using all 2P-FLIM variables. When using only τ_1_, τ_2_ and τ_avg_, it is still possible to achieve a high prediction accuracy (AUC=0.934) (Figure 3G). Next, the mean decrease accuracy and mean decrease Gini returned by the random forest model reveal that τ_1_, τ_2_ and τ_avg_ are the most important 2P-FLIM variables for macrophage classification and data segregation (Figure 3H).

### 2P-FLIM classification models are sensitive to cell heterogeneity

Macrophage polarisation heterogeneity was evaluated within each donor in response to the FCCP treatment (Figure 4A). Here, we utilised *Cell Profiler*® to evaluate and track single-cell metabolic shifts. A representative donor UMAP reveals two clusters, one dominated by IFNγ-M1 macrophages and the other dominated by IL-4-M2 macrophages (Figure 4B). Afterwards, we evaluated the classification model by plotting ROC curves and calculating its respective AUC. From Figure 4C,D it is noticeable that single-cell classification performance is affected by donor variability during the FCCP treatment. Donors D, E and A have the highest ROC-AUC values and lowest OOB errors. Finally, we measured the weight of each 2P-FLIM variable via the random forest classification model for higher ROC AUC donors, and found that τ_1_, τ_2_, τ_avg_ and α_2_ are the most important variables for classifying macrophage type at a single-cell level (Figure 4E).

## 5. Discussion

In this study, we utilise macrophage polarisation as a model system to demonstrate the effectiveness of machine learning, applied to 2P-FLIM parameters derived during subjection to small molecule metabolic perturbations to classify cell phenotype. The polarisation of human macrophages is often crudely described as two opposite phenotypes: classical activation (IFNγ-M1-Macrophages) and alternative activation (IL-4-M2-macrophages). The higher production of TNF-α in IFNγ-treated human macrophages and low IL-10 production are evidence of a macrophage classical activation(Tokunaga et al. 2018) (Figure 1A, B). While treating human macrophages with IL-4, an increase in IL-10 production and higher expression of MRC1 and CCL13, is characteristic of an alternative activation of macrophages (Artyomov et al. 2016, Martinez et al. 2006) (Figure 2D,E and F). In addition, we performed flow cytometry of polarised macrophages using antibodies to detect CD80, CD86, CD163 and CD206 surface markers, further validating the intended macrophage polarisation (supplementary Figure 2). Extending from this, extracellular flux analysis revealed a higher acidification ratio (ECAR) and a lower oxygen consumption ratio (OCR) for IFNγ-M1-macrophgaes during the different stages of metabolic inhibition when compared with IL-4-M2 macrophages (Figure 2G, J). By calculating the basal glycolysis rate and maximum glycolysis, IFNγ-M1-macrophages presented higher glycolytic rates (Figure 2H, I). The acidification (from ECAR) is linked with the production of lactate as a by-product of glycolysis, which reduces extracellular pH (Wang et al. 2018). Regarding IL-4-M2 macrophages, a reduced ECAR and increased OCR during the extracellular flux treatments (Figure 2G, J) together with a higher max respiration potential was observed when compared with untreated macrophages. However, no difference was observed at the basal respiration measures (Figure 2K, L). These results are associated with a higher dependence of OxPhos as a more active metabolic machinery in IL-4-M2 macrophages. In order to fuel the upregulation of the TCA cycle and the ETC, the mitochondria need to consume more oxygen at the complex IV site (O’Neill et al. 2016, Van den Bossche et al. 2015).

We next sought to underline 2P-FLIM as a complimentary and more advantageously, a non-invasive spatial evaluation of macrophage metabolism reflecting their induced polarisation. Recapitulating the sequence of metabolic enzyme inhibition which formed a basis of the extracellular flux analysis, sequential 2P-FLIM micrographs of IFNγ- or IL-4-polarised macrophages subjected to the same regimes were acquired (Figure 3A). A reduced τ_avg_, observed with IFNγ-M1 macrophages reflects a higher relative amount of free NAD(P)H (which has characteristic short fluorescence lifetimes), indicative with glycolysis (Perottoni et al. 2021, Walsh et al. 2013). In contrast IL-4-M2 macrophages had higher τ_avg_ - reflective of OxPhos, due to increased proportion of longer lifetime, protein-bound NAD(P)H (Okkelman et al. 2019, Walsh et al. 2020) (Figure 3B). Varying interpretations of ORR are reported in previous studies (Varone et al. 2014, Walsh et al. 2020) and for this study, we adopt a lower ORR reflecting a higher fraction of NAD(P)H and a lower FAD^+^ associated with upregulation of OxPhos (Equation 3)(Neto et al. 2020). IL-4-M2 macrophages exhibited a lower ORR across the treatments in contrast with IFNγ-M1 macrophages, validating a higher dependency of IL-4-M2 macrophages on OxPhos when compared with IFNγ-M1 macrophages. A heatmap overviewing the 2P-FLIM output variables was compiled showcasing the shifts promoted by the different treatments in both phenotype-directed human macrophages (Figure 3D). With IL-4-M2 macrophages, there is a noticeable increase in τ_1_, τ_2_ and τ_avg_ and an appreciable decrease in ORR when compared with IFNγ-M1 treated macrophages. The tending increase of NAD(P)H fluorescence lifetimes is further exacerbated after treatment with FCCP in IL-4-M2 macrophages. Indeed, given the potency of FLIM to imaging and measure NAD(P)H and FAD^+^ others have employed small molecules to gain further information about the dynamics of metabolic machinery in states of disease and differentiation. For instance; during stem cell osteogenic differentiation; Guo et al. tested oligomycin A (mitochondrial respiration inhibitor) as an experimental treatment in parallel to standard osteogenic media and noted a reduced NAD(P)H average lifetime (therefore reduced oxidative phosphorylation); lower oxygen consumption and osteogenic differentiation, and increased lactate production (Guo et al. 2015). The heterogeneous response towards metabolic inhibitors by IFNγ-M1 or IL-4-M2 macrophages yields clustering patterns in the two-dimensional projected space via the UMAP method (Figure 3E). When analysing each treatment individually, measurements emanating from the FCCP treatment yielded the highest segregation across all donors between IFNγ-M1 and IL-4-M2 macrophages (Figure 3F). Higher NAD(P)H fluorescence lifetimes such as τ_1_, τ_2_ and τ_avg_ are attributable to two major factors: increases in NADPH concentrations and microenvironmental shifts(Blacker et al. 2014, Schaefer et al. 2019). FCCP functions as an uncoupler of mitochondria inner membrane allowing unhinged proton flux to the mitochondria matrix causing a reduction of mitochondrial pH, increasing effectively the fluorescence lifetime of NAD(P)H. This compliments existing studies whereby Schaefer et al., reported an increased NAD(P)H fluorescence lifetime due to reduced mitochondria pH elicited by FCCP treatment (Blinova et al. 2005, Schaefer et al. 2017). Another impact of FCCP treatment is an increase in ETC activity indicated by increased oxygen consumption by IL-4-M2 macrophages (Figure 2J). This promotes an increase of NADH and FAD^+^ directly impacting the ORR. One would have expected the ORR to increase after the FCCP treatment as observed in IFNγ macrophages. However, for IL-4-M2 macrophages, the ORR begins to decrease. FCCP induction of maximal ETC activity increases the demand for NADH and FADH_2_ which causes a concomitant increase of the FAO and TCA cycle activity already upregulated in IL-4-M2 macrophages (Ludtmann et al. 2014). This demand results in a reduced pool of FAD^+^ and an increase of NADH, effectively reducing ORR (Akie et al. 2015, Ludtmann et al. 2014, Viola et al. 2019). The higher heterogeneity in FCCP response is due to the effect of FCCP on mitochondrial membrane and the higher dependence of FAS/FAO of IL-4-M2 macrophages basally(O’Neill et al. 2016). Blacker et al. report this in great detail where, when seeking to separate NADH and NADPH fluorescence in live cells and tissues using FLIM, they treated wild-type HEK293 cells by inhibition of mitochondrial oxidative phosphorylation using rotenone (10⍰μM) or uncoupling using FCCP (1⍰μM) ^52^. This study provides excellent insight into NADH and NADPH dynamics and separation, and the treatment of FCCP has a similar effect on HEK293 cells as the IFNγ-M1 macrophages reported in our study; whereby FCCP has less a significant impact on our IL-4-M2 macrophages. Here, FCCP uncoupling promotes higher ETC, higher TCA activity impacting the ORR and, at the same time, a decreased mitochondrial pH increases the fluorescence lifetimes of NAD(P)H (Blinova et al. 2005, Schaefer et al. 2017). Regarding IFNγ-M1 macrophages, their lower dependence on ETC, TCA, FAO and lower mitochondria membrane potential while producing most of its ATP by glycolysis, makes the impact of FCCP on the mitochondria and fluorescence lifetimes less pronounced.

Machine learning models were then adopted to study the endpoint variables of the 2P-FLIM obtained during the FCCP treatment of IFNγ-M1 macrophages and IL-4-M2 macrophages (Figure 3G, H). Here, a high ROC AUC of 0.944 was achieved when classifying IFNγ-M1 and IL-4 M2 macrophages. When classification was performed using three variables: τ_1_, τ_2_ and τ_avg_, the classification accuracy decreased slightly to 0.934 (Figure 3G). When applying random forest for feature importance ranking based on mean decrease in accuracy and mean decrease in Gini, τ_1_, τ_2_ and τ_avg_ are the best performing 2P-FLIM variables for classifying an IFNγ-M1 or an IL-4-M2 macrophage when subjected to the FCCP treatment (Figure 3H). This also implies that these FLIM variables are divergent in IFNγ-M1 and IL-4-M2 macrophages. This outcome agrees with our previous results showing that FCCP highly impacts the NAD(P)H fluorescence lifetimes in IFNγ-M1 and IL-4-M2 macrophages (figure 3D). This accuracy of 2P-FLIM to classify cell phenotype compliments the random forest machine learning models approach of Walsh et al. classifying CD3+ and CD3+CD*+ T-cell activation(Walsh et al. 2020).

Depending on the investigation being applied, shifts observed in cellular metabolism, cytokine production or gene expression are typically a cumulative output from a broad population. We investigated the clustering pattern of IFNγ-M1 macrophages and IL-4-M2 macrophages using single-cell data within individual donors (Figure 4A, B), and found that the IFNγ-M1 macrophage and IL-4-M2 macrophage appeared as distinct separate clusters within a donor. Classifying IFNγ-M1 and IL-4-M2 macrophages at a single-cell level yielded some varied results, with four donors providing acceptable predicting performance (ROC AUC>0.75) (Figure 4C, D). There is some cell-to-cell variability which could stem from the uptake capacity of FCCP and other treatments in our experiments as well the underlying health of the donors which is not available(Smiley et al. 1991, Stiebing et al. 2017). Nonetheless, for the four donors with the most superior performance during classification, τ_1_, τ_2_, τ_avg_ and α_2_ are the most potent variables highlighted by the random forest model, which agrees with the case of cell-pool analysis. One way to improve single-cell classification would be to increase the overall number of cells analysed to ensure a stronger classification.

2P-FLIM imaging has several advantages when compared with traditional metabolic assays and methods to classify and validate macrophage metabolism. It enables spatial and temporal resolution in a non-invasive manner, allowing single-cell and cell-to-cell evaluations into cellular heterogeneity in a basal and interrogated mode. It requires no fixation nor staining of cells and can be performed in real-time with only a small number of cells. In this work, we demonstrate the feasibility of using 2P-FLIM as a tool to distinguish and classify opposing human macrophage polarisation states based on cellular metabolism and fluorescence lifetimes variables. Visualisation of the data show a clear classification of IFNγ-M1 and IL-4-M2 macrophages in response to real-time imaging when treated with FCCP. The excellent performance of machine learning models, applied on the data extracted from the non-invasive technique, underlines further the efficiency of this workflow. This workflow can be easily adopted to non-invasively characterise macrophage polarisation in *in vivo models* and *in vitro* multicellular organoid models to study foreign body interactions, biomaterial assessment, pharmaceutical research and screening and clinical applications such as disease diagnosis. Precise regulation of macrophage activation state is key to understanding disease control, tissue homeostasis and implant response, with this regulation shown to be directly related with macrophage intracellular metabolism(O’Neill et al. 2016). Therefore, impaired macrophage metabolism results in impaired function such as the case of diabetes, the foreign body response to biomaterials, obesity or cancer(Mantovani et al. 2010, McNelis et al. 2014).

## 6. Acknowledgements

NN is supported by a Trinity College Dublin, Provost’s PhD Award and the TCD FLIM core unit directed by MM is supported by a SFI Infrastructure Programme: Category D Opportunistic Funds Call (16/RI/3403). This work was also partially supported by EPSRC and SFI Centre for Doctoral Training in Engineered Tissues for Discovery, Industry and Medicine, Grant Number EP/S02347X/1. This publication has emanated from research supported in part by a grant from Science Foundation Ireland (SFI) and the European Regional Development Fund (ERDF) under grant number 13/RC/2073_P2.

## 7. Author Contributions

NGBN and MGM conceptualised the study. NGBN, SOR, MGM designed experiments and wrote the paper. NGBN, SOR, MZ, HKF performed experiments, collected and analysed the data. MZ, AD, MGM revised the manuscript.

## 8. Competing interests

The authors declare no competing interests.

## 9. Materials & Correspondence

Correspondence to Michael G. Monaghan

## Notes

### Competing Interest Statement

The authors have declared no competing interest.

