## Supplementary File for "Non-Invasive classification of macrophage polarisation by 2P-FLIM and machine learning"

Supplementary Table 1 – *mtry*, *ntree* and OOB hyperparameters of random forest model

| **Donor** | **mtry** | **ntree** | **OOB (%)** |
|---|---|---|---|
| All (Global data) | 100 | 2 | 16.67 |
| A | 400 | 5 | 23.53 |
| B | 200 | 3 | 38.06 |
| C | 100 | 2 | 30.17 |
| D | 300 | 4 | 10.05 |
| E | 150 | 2 | 19.55 |
| F | 150 | 3 | 28.77 |

Supplementary Table 2 – Calculation of basal glysolysis, max glycolycic, basal respiration, max respiration for ECAR/OCR experimental setup.

| Rate | Calculation |
|---|---|
| **Basal glycolysis** | Average ECAR values prior to oligomycin treatment – non-glycolytic ECAR |
| **Max glycolysis** | Average ECAR values after oligomycin and before FCCP treatment |
| **Basal respiration** | Average OCR values prior to oligomycin treatment – nonmitochondrial OCR |
| **Max respiration** | Average OCR values after FCCP and before rotenone/antimycin A treatment |

Supplementary Figure 1 – Correlation matrix of 2P-FLIM NAD(P)H variables

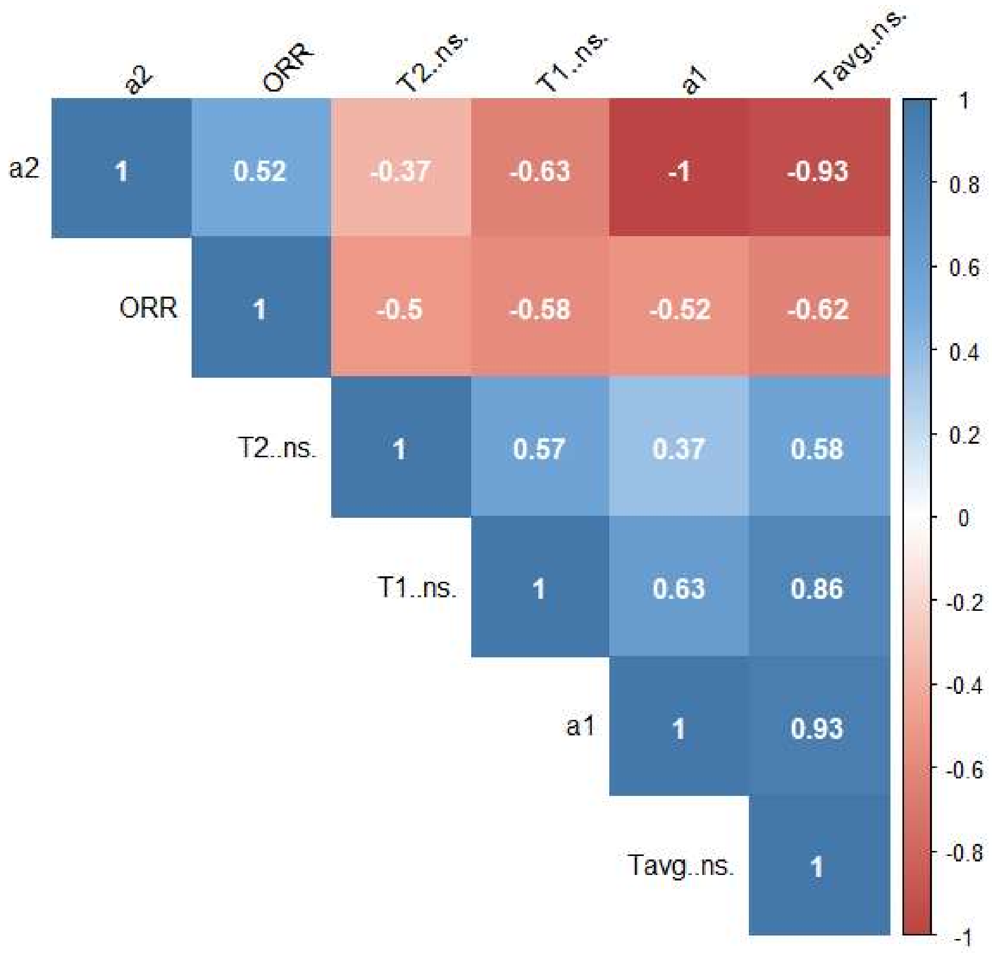

Supplementary Figure 2 – Validation of macrophage polarisation using Flow Cytometry

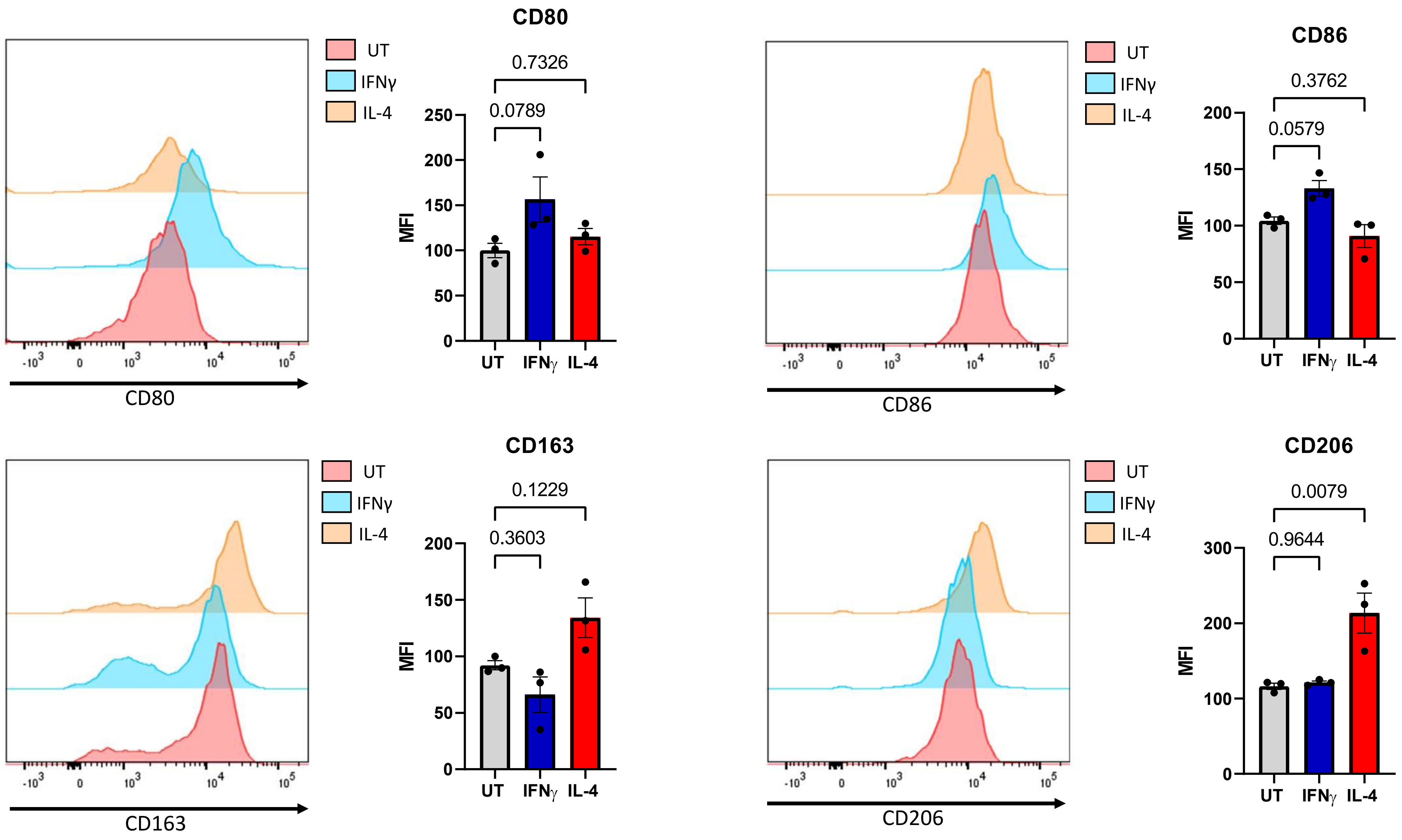
